# FGIN-1-27, an agonist at translocator protein 18 kDa (TSPO), produces anti-anxiety and anti-panic effects in non-mammalian models

**DOI:** 10.1101/056507

**Authors:** Monica Gomes Lima, Jonathan Cueto-Escobedo, Juan Francisco Rodríguez-Landa, Caio Maximino

## Abstract

FGIN-1-27 is an agonist at the translocator protein 18 kDa (TSPO), a cholesterol transporter that is associated with neurosteroidogenesis. This protein has been identified as a peripheral binding site for benzodiazepines; in anamniotes, however, a second TSPO isoform that is absent in amniotes has been implicated in erythropoiesis. Functional conservation of the central benzodiazepine-binding site located in the GABA_A_ receptors has been demonstrated in anamniotes and amniotes alike; however, it was not previously demonstrated for TSPO. The present investigation explored the behavioral effects of FGIN-1-27 on an anxiety test in zebrafish (*Danio rerio*, Family: Cyprinide) and on a mixed anxiety/panic test on wall lizards (*Tropidurus oreadicus*, Family: Tropiduridae). Results showed that FGIN-1-27 reduced anxiety-like behavior in the zebrafish light/dark preference test similar to diazepam, but with fewer sedative effects. Similarly, FGIN-1-27 also reduced anxiety- and fear-like behaviors in the defense test battery in wall lizards, again producing fewer sedative-like effects than diazepam; the benzodiazepine was also unable to reduce fear-like behaviors in this species. These results A) underline the functional conservation of TSPO in defensive behavior in anamniotes; B) strengthen the proposal of using anamniote behavior as models in behavioral pharmacology; and C) suggest TSPO/neurosteroidogenesis as a target in treating anxiety disorders.

## 1. Introduction

The translocator protein 18 kDa (TSPO, mitochondrial benzodiazepine receptor [MBR], peripheral benzodiazepine receptor) was first identified as a peripheral binding site for diazepam, but later identified as part of the mitochondrial cholesterol transport pathway that is associated with the regulation of cellular proliferation, immunomodulation, porphyrin transport and heme biosynthesis, anion transport, regulation of steroidogenesis, and apoptosis (Casellas et al., 2002). This transporter is highly expressed in steroidogenic tissues. In the central nervous system, its expression is mainly restricted to ependymal cells and glia, in which it is responsible for the local synthesis of neuroactive steroids such as allopregnanolone (Papadopoulos et al., 2006). This latter neurosteroid, in its turn, positively modulates GABA_A_ receptors, especially those involved in tonic inhibition (Smith et al., 2009; Maguire et al., 2012), producing anxiolytic-(Fernández-Guasti and Picazo, 1995; Schüle et al., 2014) and antidepressant-like effects (Khisti et al., 2000; Rodríguez-Landa et al., 2007, 2009), and modulates panic attacks (Bali and Jaggi, 2014; Lovick, 2014). As such, TSPO has been proposed as a pharmacological target for the treatment of neurological and psychiatric disorders associated with decreased GABAergic tone, such as anxiety disorders (Romeo et al., 1993; de Mateos-Verchere et al., 1998; Kita et al., 2004; Costa et al., 2011; Matsuda et al., 2011; Nin et al., 2011; Pinna and Rasmusson, 2012; Pinna and Rasmusson, 2012; Perna et al., 2014) and epilepsy (Ugale et al., 2004), as well as for the fine control of stress responses (Gunn et al., 2011; Maguire et al., 2012; Maguire, 2014). TSPO agonists produce anti-anxiety and anti-conflict effects in rodents with both systemic (Kita et al., 2004; Costa et al., 2011) and intra-hippocampal (Bitran et al. 2000) injections; these effects are blocked by GABA_A_ receptor antagonists (i.e. picrotoxin) and/or 5α-reductase blockers (i.e. 4-MA) and TSPO antagonists (PK 11195), implicating neurosteroidogenesis and chloride ion channel at the GABA_A_ receptors in these responses (Bitran et al. 2000). Additionally, octadecaneuropeptide, a diazepam-binding inhibitor peptide that acts through both the central benzodiazepine receptor (CBR) and TSPO produces anxiety-like behavior in mice and rats (de Mateos-Verchere et al., 1998), as well as in goldfish (Matsuda et al., 2011). These effects are spared in adrenalectomized and castrated animals, suggesting that they are not mediated by peripheral steroidogenesis, but rather by the production of neurosteroids in the brain (Romeo et al., 1993).

TSPO is highly conserved, being present in Bacteria, Archaea and Eukarya domains (Fan and Papadopoulos, 2013). Anamniotes and invertebrates possess a single isoform, while amniotes possess two TSPO isoforms (Fan et al., 2009). Interestingly, while no functional divergence is predicted to appear between *tspo* (found in invertebrates and basal vertebrates) and *tspo1* (found in amniotes), a functional divergence was detected in TSPO2 (Fan and Papadopoulos, 2013). Some of the neurobehavioral functions of this protein, on the other hand, seem to be conserved. In zebrafish, for example, benzodiazepines have been shown to affect a plethora of anxiety-like behaviors, from bottom-dwelling (Bencan et al., 2009; Egan et al., 2009) and dark preference (Maximino et al., 2010, 2011) to shoal cohesion (Gebauer et al., 2011) and cocaine withdrawal-induced anxiety-like behavior (López-Patiño et al., 2008). Likewise, benzodiazepines decrease tonic immobility duration and the following freezing and explosive behavior in a defensive behavior battery in the wall lizard (*Tropidurus oreadicus*), and also increase exploratory behavior in the same test (Maximino et al., 2014). In the separation stress paradigm, benzodiazepines attenuate separation stress-induced distress vocalizations in chicks in the anxiety phase, but not in the depression phase (Warnick et al., 2009). Thus, agonists at the CBR decrease fear- and anxiety-like behavior in both amniotes and anamniotes. Moreover, some evidence regarding the neurosteroidogenesis pathway in behavioral control has been suggested by the observation that allopregnanolone has an anticonvulsant effect in zebrafish (Baxendale et al. 2012), and that chronic fluoxetine treatment upregulates the expression of genes from the neurosteroidogenesis pathway in this species (Wong et al. 2013). These results suggest that some downstream effectors of neurosteroidogenesis are conserved, although little is known about the role of TSPO in behavioral control *per se*. A comparative approach could untangle this question, especially if species at the base of the amniote and anamniote clades are used. In this paper, we describe the behavioral effects of FGIN-1-27, a TSPO agonist, in zebrafish (*Danio rerio* Hamilton 1822, Family: Cyprinidae) and wall lizards (*Tropidurus oreadicus* Rodrigues, Family: Tropiduridae) and compare these responses with the effects of diazepam, an agonist at the CBR with tested anxiolytic and anticonvulsant actions at preclinical and clinical research.

## 2. Experimental procedures

### 2.1 Experiment 1: Effects of FGIN-1-27 and diazepam on dark preference in zebrafish

#### 2.1.1. Animals and husbandry

120 adult zebrafish from the *longfin* phenotype were acquired in a local aquarium shop and kept in collective tanks at the laboratory for at least 2 weeks before experiments started. Conditions in the maintenance tank were kept stable, as described by Lawrence (2007). Recommendations in the Brazilian legislation (Conselho Nacional de Controle de Experimentação Animal - CONCEA, 2017) were followed to ensure ethical principles in animal care and throughout experiments. This manuscript is a complete report of all the studies performed to test the effect of diazepam or FGIN-1-27 on anxiety-like behavior in zebrafish. We report how the sample size was determined, all data exclusions (if any), all manipulations, and all measures in the study.

#### 2.1.2. Drug administration

Diazepam was dissolved in 40 % propylene glycol, 10 % ethyl alcohol, 5 % sodium benzoate, and 1.5 % benzyl alcohol (Maximino et al. 2010). FGIN-1-27 was dissolved in 1% DMSO to which one or two drops of Tween 80 was added before sonication into a fine suspension (Auta et al. 1993). Drugs were diluted to their final concentrations and injected *i.p*. in a volume of 1 µL/0.1 g b.w. (Kinkel et al. 2010). For both diazepam and FGIN-1-27, the doses used were 0.14, 0.28, 0.57, 1.1, and 2.3 mg/kg. The three highest doses were chosen based on reported effects of diazepam on zebrafish (Maximino et al. 2010) or wall lizard (Maximino et al. 2014) defensive behavior, and were identical for both drugs to facilitate comparison; the lower doses were chosen in order to define a minimal effect. All drugs were acquired from Tocris Bioscience (UK).

#### 2.1.3. Experimental groups and dosage

The adult zebrafish were assigned to 12 independent groups (n= 10 fish per group): a vehicle group (propylene glycol:ethyl alcohol:sodium benzoate:benzyl alcohol for the diazepam controls, DSMO for the FGIN-1-27 controls) that received 1 µL/0.1 g b.w.; and 10 additional groups that received the doses described above for diazepam (5 groups) or FGIN-1-27 (5 groups). Sample sizes were based on Maximino *et al.* (2010). All drugs were *i.p.* injected in a volume of 1 µL/0.1 g b.w., 30 min before of the behavioral test. Animals were randomly drawn from the holding tank immediately before injection, and the order with which doses were tested was randomized *via* generation of random numbers using the randomization tool in http://www.randomization.com/. Experimenters were blinded to treatment by using coded vials for drugs. The data analyst was blinded to phenotype by using coding to reflect treatments in the resulting datasets; after analysis, data was unblinded.

#### 2.1.4. Scototaxis assay

The light/dark preference (scototaxis) assay was performed as described by Maximino *et al.*, (2013). Briefly, 30 min after injection animals were transferred individually to the central compartment of a black/white tank (15 cm height X 10 cm width X 45 cm length) for a 3-min acclimation period, after which, the doors which delimit this compartment were removed and the animal was allowed to freely explore the apparatus for 15 min. While the whole experimental tank was illuminated from above by an homogeneous light source, due to the reflectivity of the apparatus walls and floor average illumination (measured just above the water line) above the black compartment was 225 ± 64.2 (mean ± S.D.) lux, while in the white compartment it was 307 ± 96.7 lux. The following variables were recorded:

> *time spent on the white compartment*: the time spent in the white half of the tank (percentage of the trial);
>
> *squares crossed*: the number of 10 cm^2^ squares crossed by the animal in the white compartment;
>
> *latency to white compartment*: the time in seconds (s) to first entry in the white compartment; *erratic swimming:* defined as the number of zig-zag, fast, and unpredictable swimming behavior of short duration;
>
> *freezing*: the proportional duration of freezing events (in % of time in the white compartment), defined as complete cessation of movements with the exception of eye and operculum movements;
>
> *thigmotaxis*: the proportional duration of thigmotaxis events (in % of time in the white compartment), defined as swimming in a distance of 2 cm or less from the white compartment’s walls;
>
> *Number of risk assessment*: defined as a fast (<1 s) entry in the white compartment followed by re-entry in the black compartment, or as a partial entry in the white compartment (i.e., the pectoral fin does not cross the midline);

A digital video camera (Samsung ES68, Carl Zeiss lens) was installed above the apparatus to record the behavioral activity of the zebrafish. Two independent observers, blinded to treatment, manually measured the behavioral variables using X-Plo-Rat 2005 (https://github.com/lanec-unifesspa/x-plo-rat). Inter-observer reliability was at least > 0.85.

#### 2.1.5. Statistical analysis

The data were analyzed using two-way ANOVAs, with drug and dose as between-subjects factors. Values of *p* ≤ 0.05 in the ANOVA were followed by Tukey’s HSD whenever appropriate; planned comparisons were between different doses of a given drug and its vehicle and between same doses of both drugs. All statistical analyses were made using R 3.1.3. Data are presented graphically as individual points superimposed over boxplots representing medians and interquartile ranges with Tukey whiskers. To address the question of normality, graphs also present the mean of each group, connected by lines.

#### 2.1.6. Data availability

Full data can be found in our GitHub repository (https://github.com/lanec-unifesspa/fgin-1-27/tree/master/zebrafish)

### 2.2. Experiment 2: Effects of FGIN-1-27 and diazepam on the defense test battery in *Tropidurus oreadicus*

#### 2.2.1. Animals and husbandry

120 adult wall lizards (*Tropidurus oreadicus*) of either sex, ranging from 61-96 mm in rostro-cloacal length, were captured in Marabá/PA, Brazil, between February and March 2015. The animals were inspected for mites, which were removed with forceps before treatment with demiting solution as described by the manufacturer (Reptile Relief, Natural Chemistry, Norwalk, USA). All of the lizards were treated with 50 mg/kg fenbendazole (Vetnil, Brazil), p.o., and then housed according to recommendations for anoline lizards (Sanger et al. 2008) for at least 2 weeks before the experiments began. Animals were housed in groups of four in Plexiglas standard laboratory cages (42 cm length x 27.5 cm width x 21 cm height) with mango tree sticks collected from the outdoors to provide perches. Before using the sticks, they were sterilized for 15 min in an autoclave. To prevent escape, screen meshes were inserted in the cage tops. The bottoms of the cages were covered with synthetic cage carpet (Repti Cage Carpet CC-10, Zoo Med, Costa Mesa, USA) placed above a heater plate (Repti Therm U.T.H. Under Tank RH-6, Zoo Med, Costa Mesa, USA) that kept the temperature above the carpet at an average of 28°C. The cages were misted with water twice daily, thus raising the humidity within each cage to approximately 85% (Sanger et al. 2008). The animals had *ad libitum* access to drinking water. The animals were fed three times weekly with commercial ration (Shrimp mix, Nutral, Monte Mor, Mexico) and once per week with captured crickets. In the absence of Brazilian guidelines for lizards, the guidelines for the husbandry of *Anolis* lizards by Sanger *et al.* (2008) were used, in an attempt to ensure ethical principles in animal care throughout experiments.

#### 2.2.2. Drug administration

FGIN-1-27 and diazepam were prepared as in Experiment 1 and injected intraperitoneally in a volume of 0.5 ml of either vehicle or drug 30 min before behavioral tests.

#### 2.2.3. Experimental groups and dosage

Adult lizards were assigned to 12 independent groups (n= 10 lizards per group): a vehicle group (propylene glycol:ethyl alcohol:sodium benzoate:benzyl alcohol for the diazepam controls, DSMO for the FGIN-1-27 controls) that received 1 µL/0.1 g b.w.; and 10 additional groups that received the doses described above for diazepam (5 groups) or FGIN-1-27 (5 groups). Sample sizes were based on Maximino *et al.* (2014). All drugs were *i.p*. injected in a volume of 0.5 ml per animal, 30 min before of the behavioral test. Animals were randomly drawn from the cage immediately before injection, and the order with which doses were tested was randomized *via* generation of random numbers using the randomization tool in http://www.randomization.com/. Experimenters were blinded to treatment by using coded vials for drugs. The data analyst was blinded to phenotype by using coding to reflect treatments in the resulting datasets; after analysis, data was unblinded.

#### 2.2.4. Defense test battery

The defense test battery was applied as described by Maximino *et al.* (2014). Briefly, tonic immobility was induced by carefully placing the animal on its back in the center of a 10 cm diameter circular open field and applying mild pressure to the thorax and pelvis while restraining the limbs. When the lizard ceased struggling, it was slowly released, and the time taken for it to resume an upright posture was recorded (Hennig 1979). After the animal spontaneously ceased tonic immobility (TI), the following behavioral endpoints were recorded after each of these manipulations:

> *tonic immobility (TI)*: the total duration, in minutes (min), in a rigid supine posture after release;
>
> *freezing*: the lack of limb, neck, or tongue movements for more than 5 s in an upright position;
>
> *circling*: a high-velocity escape attempt with a latency of less than 10 s after release, usually leading to circling around the edges of the apparatus, and quantified as the number of complete circles made near the walls;
>
> *tongue-flicking*: repeatedly licking the air with the tongue;
>
> *ventilatory frequency*: the average number of inspiratory responses per minute;
>
> *total locomotion*: the number of 2 cm^2^ squares crossed by normal locomotor responses (i.e., not concomitant to circling); can be superimposed to tongue-flicking.

According to a previous work, tonic immobility, circling, ventilatory frequency, and freezing behaviors are considered responses to fear since panicolytic drugs are able to reduce its levels; tongue-flicking is considered an exploratory behavior; and total locomotion signals hyper- or hypolocomotor/sedative effects (Maximino et al., 2014).

A digital video camera (Samsung ES68, Carl Zeiss lens) was installed above the apparatus to record the behavioral activity of the lizards. Two independent observers, blinded to treatment, manually measured the behavioral variables using EthoLog 2.2 (Ottoni 2000). Inter-observer reliability was at least > 0.85.

#### 2.2.5. Statistical analysis

The data were analyzed using two-way ANOVAs, with drug and dose as between-subjects factors. Values of *p* ≤ 0.05 in the ANOVA were followed by Tukey’s HSD whenever appropriate; planned comparisons were between different doses of a given drug and its vehicle and between same doses of both drugs. All statistical analyses were made using R 3.1.3. Data are presented graphically as individual points superimposed over boxplots representing medians and interquartile ranges with Tukey whiskers. To address the question of normality, graphs also present the mean of each group, connected by lines.

#### 2.2.6. Data availability

Full data can be found in our GitHub repository (https://github.com/lanec-unifesspa/fgin-1-27/tree/master/lizards)

## 3. Results

### 3.1. Experiment 1

Main effects of drug (F_1, 107_ = 91.216, p < 0.0001) and dose (F_5, 107_ = 32.173, p < 0.0001), as well as an interaction effect (F_5,107_ = 4.939, p = 0.000416) were observed in time spent into the white compartment (Figure 1A). Post-hoc tests uncovered differences between all doses except the highest in relation to controls in diazepam-treated animals (p < 0.001) and between all doses in relation to controls in FGIN-1-27-treated animals (p < 0.001). Finally, differences were also observed between FGIN-1-27- and diazepam-treated animals at doses of 0.57 mg/kg (p = 0.0012), 1.1 mg/kg (p < 0.0001), and 2.3 mg/kg (p = 0.0002).

**Figure 1.**
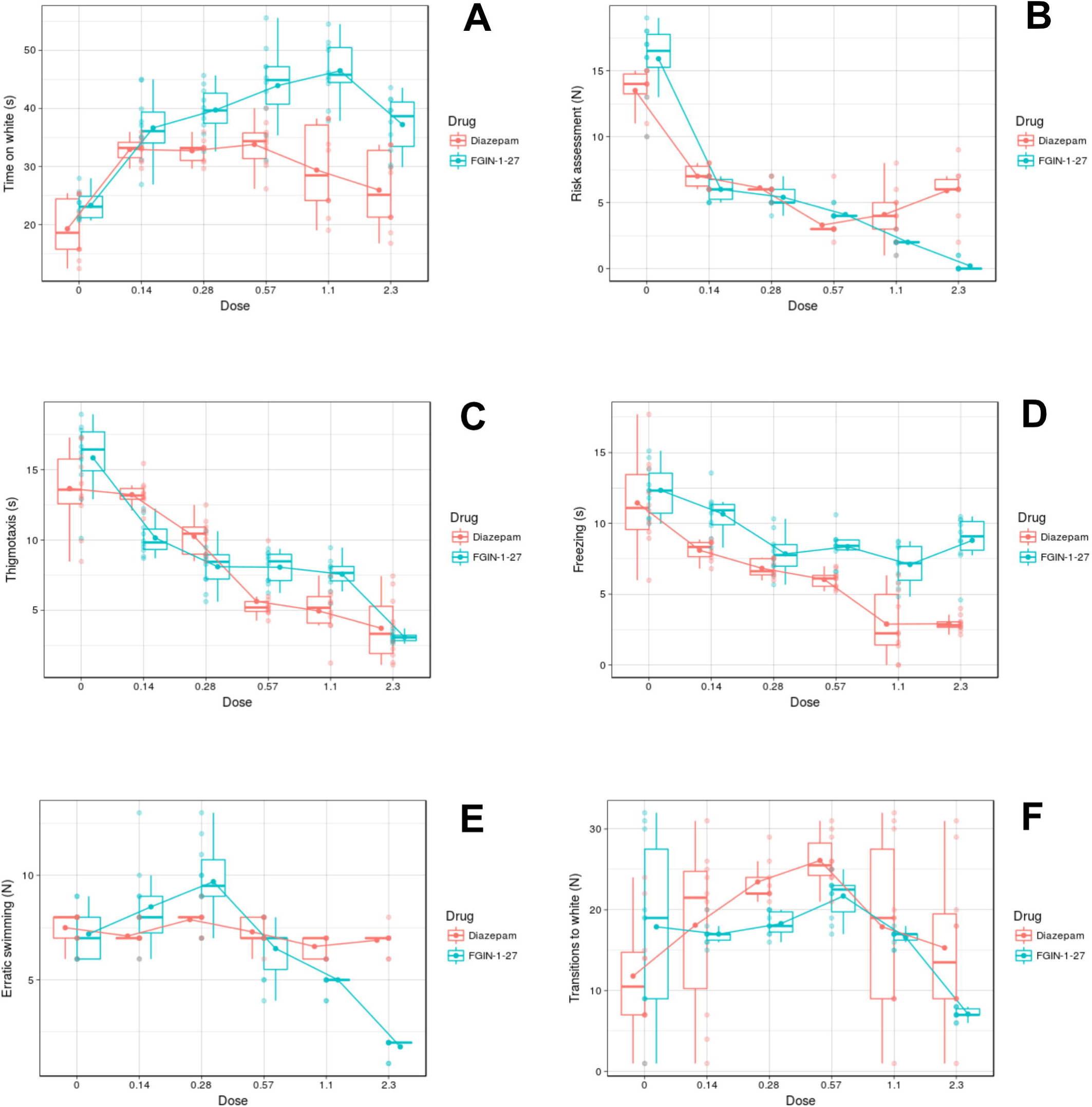
Effects of FGIN-1-27 (blue) and diazepam (pink) on (A) scototaxis, (B) risk assessment, (C) thigmotaxis, (D) freezing, (E) erratic swimming, and (F) total locomotion in the light/dark test in zebrafish (*Danio rerio*). Data are presented as individual points (pale dots) superimposed over boxplots representing medians and interquartile ranges with Tukey whiskers. To address the question of normality, graphs also present the mean of each group (solid dot), connected by lines.

Main effects of drug (F_1, 107_ = 17.64, p < 0.0001) and dose (F_5, 107_ = 207.77, p < 0.0001), as well as an interaction effect (F_5, 107_ = 20.11, p < 0.0001), were found for risk assessment (Figure 1B). Post-hoc tests uncovered differences between vehicle-treated and diazepam-treated animals at all doses (p < 0.0001), and between vehicle-treated and FGIN-1-27-treated animals at all doses (p < 0.0001). Moreover, differences between diazepam- and FGIN-1-27-treated animals were observed at 1.1 mg/kg (p = 0.012) and 2.3 mg/kg (p < 0.0001).

A main effect of dose (F_5, 107_ = 111.547, p < 0.0001), but not drug (F_1, 107_ = 0.643, p = 0.424), was found for thigmotaxis (Figure 1C), and an interaction effect was also found (F_5, 107_ = 10.536, p < 0.0001). Post-hoc tests unveiled differences between vehicle-treated and diazepam-treated animals at doses above 0.28 mg/kg (p < 0.01), as well as between all FGIN-1-27 doses and vehicle-treated animals (p < 0.0001). Differences were also observed between diazepam- and FGIN-1-27-treated zebrafish at doses of 0.14 mg/kg (p = 0.0065) and 1.1 mg/kg (p = 0.044).

Main effects of drug (F_1, 107_ = 89.134, p < 0.0001) and dose (F_5, 107_ = 47.418, p < 0.0001), as well as an interaction effect (F_5. 107_ = 6.921, p < 0.0001), were found for freezing (Figure 1D). Post-hoc tests uncovered differences between vehicle-treated and diazepam-treated animals at all doses (p < 0.001), and between FGIN-1-27-treated and vehicle-treated animals at 0.28 mg/kg and higher doses (p < 0.001). Differences were also found between diazepam- and FGIN-1-27-treated zebrafish at doses of 0.14 mg/kg (p = 0.03), 1.1 mg/kg (p < 0.0001), and 2.3 mg/kg (p < 0.0001).

Main effects of drug (F_1, 107_ = 14.57, p = 0.000226) and dose (F_5, 107_ = 42.64, p < 0.0001), as well as an interaction effect (F_5, 107_ = 26.48, p < 0.0001), were found for erratic swimming (Figure 1E). Post-tests uncovered no statistically significant differences between vehicle-treated and diazepam-treated animals at any dose (p > 0.05), while FGIN-1-27-treated animals were significantly different between vehicle-treated animals at doses of 0.28 mg/kg (p = 0.0001), 1.1 mg/kg (p < 0.001) and 2.3 mg/kg (p < 0.0001). Also, differences were observed between FGIN-1-27- and diazepam-treated animals at 0.28 mg/kg (p = 0.021), 1.1 mg/kg (p = 0.05), and 2.3 mg/kg (p < 0.0001).

Finally, no main effects of drug (F_1, 107_ = 3.526, p = 0.0631), but a main dose effect (F_5, 107_ = 9.088, p < 0.0001) as well as a suggestive interaction effect (F_5, 107_ = 2.778, p = 0.0212) were found for number of transitions into the white compartment (Figure 1F). Post-hoc tests uncovered differences between vehicle- and diazepam-treated zebrafish at doses of 0.28 mg/kg and 0.57 mg/kg (p < 0.01) and vehicle- and FGIN-1-27-treated zebrafish at the highest dose (p = 0.003). No differences were found between diazepam- and FGIN-1-27-treated animals.

### 3.2. Experiment 2

Main effects of drug (F_1, 108_ = 248.5, p < 0.0001) and dose (F_1, 108_ = 93.39, p < 0.0001), as well as an interaction effect (F_1, 108_ = 60.09, p < 0.0001) were found for TI duration (Figure 2A). Post-hoc tests found statistically significant differences between vehicle- and diazepam-treated lizards at doses of 0.28-1.1 mg/kg (p < 0.05) and vehicle- and FGIN-1-27-treated lizards at all doses except the highest (p < 0.0001). Differences between FGIN-1-27- and diazepam-treated lizards were observed at all doses except the higher (p < 0.0001).

**Figure 2.**
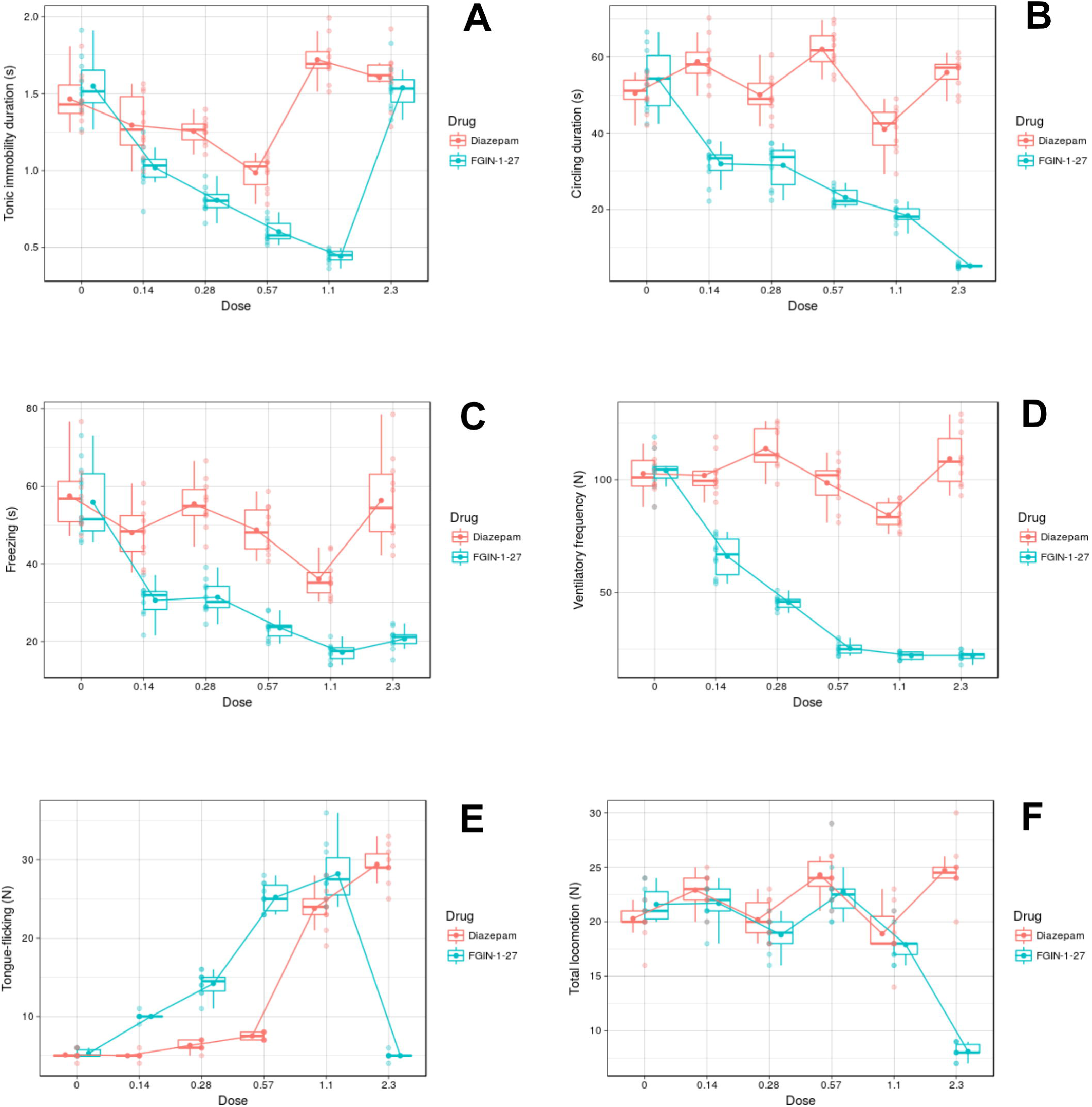
Effects of FGIN-1-27 (blue) and diazepam (pink) on (A) tonic immobility, (B) circling responses, (C) freezing, (D) ventilatory frequency, (E) tongue-flicking, and (F) total locomotion on wall lizards (*Tropidurus oreadicus*). Data are presented as individual points (pale dots) superimposed over boxplots representing medians and interquartile ranges with Tukey whiskers. To address the question of normality, graphs also present the mean of each group (solid dot), connected by lines.

Main effects of drug (F_1, 108_ = 771.29, p < 0.0001) and dose (F_5, 108_ = 59.58, p < 0.0001), as well as an interaction effect (F_5, 108_ = 66.98, p < 0.0001) were found for circling behavior (Figure 2B). Post-hoc tests revealed differences between vehicle- and FGIN-1-27 treated lizards at all doses (p < 0.0001), and between vehicle- and diazepam-treated animals at 1.1 and 2.3 mg/kg (p < 0.05). Differences between diazepam- and FGIN-1-27-treated lizards were also found for all doses (p < 0.0001).

Main effects of drug (F_1. 108_ = 291.58, p < 0.0001) and dose (F_5, 108_ = 45.09, p < 0.0001), as well as an interaction effect (F_5, 108_ = 14.63, p < 0.0001) were found for freezing (Figure 2C). Post-hoc tests identified significant differences between vehicle- and diazepam-treated animals at 1.1 mg/kg (p < 0.0001) and between vehicle- and FGIN-1-27-treated animals at all doses (p < 0.0001). Differences between diazepam- and FGIN-1-27-treated lizards were also found for all doses (p < 0.0001). A drug main effect (F_1, 108_ = 1487.5, p < 0.0001) and a dose main effect (F_5, 108_ = 110.54, p < 0.0001), as well as an interaction effect (F_5. 108_ = 86.71, p < 0.0001) were observed for ventilatory frequency (Figure 2D). Post-hoc tests uncovered differences between vehicle- and diazepam-treated animals at 1.1 mg/kg (p < 0.0001), and between vehicle- and FGIN-1-27-treated animals at all doses (p < 0.0001). Finally, statistically significant differences were found between diazepam- and FGIN-1-27-treated animals at all doses (p < 0.0001).

Main effects of drug (F_1, 108_ = 34.26, p < 0.0001) and dose (F_5, 108_ = 413.3, p < 0.0001), as well as interaction effect (F_5, 108_ = 351.08, p < 0.0001) were found for tongue-flicking (Figure 2E). Post-hoc tests revealed significant differences between vehicle- and diazepam-treated lizards at doses of 1.1 and 2.3 mg/kg (p < 0.0001), and between vehicle- and FGIN-1-27-treated lizards at all doses except the highest (p < 0.0001). Moreover, statistically significant differences between diazepam- and FGIN-1-27-treated lizards at all evaluated doses were found (p < 0.0001).

Main effects of both drug (F_1, 108_ = 91.53, p < 0.0001) and dose (F_5, 108_ = 36.28, p < 0.0001), as well as an interaction effect (F_5, 108_ = 56.62, p < 0.0001) were found for total locomotion (Figure 2F). Significant differences were found between vehicle- and diazepam-treated lizards at 0.57 and 2.3 mg/kg (p < 0.0003) and vehicle- and FGIN-1-27-treated lizards at 1.1 and 2.3 mg/kg (p < 0.002). Finally, statistically significant differences were also found between diazepam- and FGIN-1-27-treated animals at 2.3 mg/kg (p < 0.0001).

## 4. Discussion

In the present work we demonstrated that the TSPO agonist FGIN-1-27 produced a decrease in defensive behaviors in both wall lizards and zebrafish. Results can be summarized as follows: a) in zebrafish, FGIN-1-27 decreased anxiety-like behaviors including scototaxis, thigmotaxis, freezing and risk assessments. B) In wall lizards FGIN-1-27 decreased defensive behaviors as tonic immobility, ventilatory frequency, freezing and circling, while increasing tongue-flicking. C) The pharmacological sensitivity varied between species, in which the anxiolytic profile of FGIN-1-27 was similar to diazepam in zebrafish while in lizards FGIN-1-27 seemed to be more effective than diazepam.

### 4.1. Defense test battery

In the lizard defense test battery, induction of TI produces a stereotypical behavioral pattern in which is followed by either freezing or “explosive” circling behavior; after that, the animal emits exploratory behavior, marked by thigmotaxis (“wall-hugging”) and tongue-flicking (Maximino e*t al*., 2014). This sequence resembles defensive behavior at increasing predatory imminence continua (Fanselow and Lester, 1988) – that is, defensive behavior in this test shifts from panic-like, circa-strike behavior towards sustained risk assessment (i.e., tongue-flicking, thigmotaxis). Moreover, in a previous experiment panicolytic drugs (i.e., alprazolam, imipramine) decreased TI duration, freezing and circling behavior; while diazepam (0.5 mg/kg) increased tongue-flicking and decreased thigmotaxis (Maximino *et al*., 2014). In the present experiment, FGIN-1-27 produced a wider range of effects than diazepam: the TSPO agonist affected exploratory behavior (consistent with an anxiolytic-like effect), TI duration and post-TI behavior as circling and freezing (consistent with a panicolytic effect). Diazepam, on the other hand, did not modify tonic immobility, circling and freezing and only affected exploratory behavior (tongue flicking behavior) while it had a hormetic (inverted-U shape) profile on TI duration. It was previously suggested that TI and post TI behaviors can be affected by panicolytic drugs but not anxiolytic drugs such as diazepam (Maximino et al., 2014). On the other hand, FGIN-1-27 modified these behaviors. Differences in the effects between diazepam and FGIN-1-27 can be expected since FGIN-1-27 binds to TSPO/MDR and does not bind to the CBR at GABA_A_ receptors (Romeo et al., 1992), which mediate the anxiolytic effects of diazepam (Enna, 2007), giving to FGIN-1-27 a different pharmacological profile; a hypothesis that require specific protocols to be experimentally explored.

### 4.2. Dark preference test

A complementary profile was observed in the zebrafish scototaxis assay, which confirm the actions of FGIN1-27 modulating emotional states by decreasing defensive behaviors triggered by stress. In these subjects, diazepam produced an inverted-U shape effect on scototaxis and entries into the white compartment, monotonically decreasing freezing, thigmotaxis and risk assessment. Across multiple drug treatments (Maximino et al., 2014), thigmotaxis, risk assessment, and scototaxis are consistently affected by anxiolytic or anxiogenic drugs. Anxiolytic drugs, such as benzodiazepines, produce a hormetic profile in several behavioral tests in zebrafish (Bencan et al., 2009; Cachat et al., 2010; Sackerman et al., 2010; Vada et al., 2015). Previous experiments also demonstrated that diazepam was anxiolytic at 1.25 mg/kg, but not 2.5 mg/kg, in the scototaxis assay (Maximino *et al*., 2011). These hormetic effects in fish resemble those observed in mammals treated with diazepam or flavonoids that interact with the GABA_A_ receptors (File and Pelow, 1985; Auta et al., 1993; Carro-Juarez et al., 2012) and also in the defense test of lizard of the present experiment. This effect has been presumably associated to sedative actions (Auta et al., 1993), which coincides with our findings that diazepam and FGIN-1-27 produced anxiolytic-like effects, with locomotor-impairing effects at the highest dose. Although anxiolytic-like and locomotor effects differ between diazepam and FGIN-1-27 in this work, contrast in the pharmacological profile (i.e. anxiolytic, anticonvulsant, and sedative) between these drugs has been previously reported (Okuyama et al., 1999; Auta el al., 2003). It has been suggested that in addition to the effects on TSPO, FGIN-1-27 also acts indirectly on the GABA_A_ receptor by facilitating production of neurosteroids that bind to the receptor. Neurosteroids have an specific binding domain in GABA_A_ receptor, through which they can produce anxiolytic effects (Enna, 2007), whereas diazepam binds to the benzodiazepine binding domain of GABA (that is, the CBR). In brief, although FGIN-1-27 and diazepam might act on the same receptor they do it through different binding domains (Romeo et al., 1992), which could explain the minute variations in the effects of diazepam and FGIN-1-27 in zebrafish. For example, in rodents the anxiolytic effects of benzodiazepines, but not of FGIN-1-27, are antagonized by flumazenil (Auta et al., 1993). Additionally, there is still the possibility that neuroesteroids increased trough FGIN could be acting through a different system than the GABAergic, a possibility that remains to be explored.

### 4.3. Role of TSPO

Little is known about the role of TSPO in behavioral control in anamniotes. In goldfish (*Carassius auratus* Linn, Cyprinidae), octadecaneuropeptide increased the latency to enter a white compartment in a light/dark box, suggesting an anxiogenic-like effect (Matsuda et al., 2011). While octadecaneuropeptide is an endozepine that acts as an agonist at TSPO (Papadopoulos et al., 1991), it also acts as an antagonist at the CBR (Ferrero et al., 1986); consistently with the hypothesis that the anxiogenic-like effect of octadecaneuropeptide is mediated by the CBR, flumazenil, but not a metabotropic endozepine receptor antagonist, blocked the scototaxis effects in the goldfish (Matsuda et al. 2011). Overall, the present results suggest that drugs targeting the CBR and TSPO/MBR exert anti-anxiety effects in anamniotes, while drugs acting at the TSPO/MBR also exert anti-panic effects. It is plausible that some of the effects of diazepam were mediated by the TSPO/MBR; further experiments are needed to untangle the precise mechanisms. In this sense, the differences between actions points out differences in pharmacological sensitivity across species that contribute to understand the possible implications of using anamniote behavior as models in behavior pharmacology. Thus, while the present results suggest a functional conservation of TSPO that is concomitant to gene duplication, it is not known whether FGIN-1-27 (or even diazepam) produced its behavioral effects in lizards and zebrafish by acting on the (conserved) *tspo1* or whether some effects are also mediated by *tspo*. Further experiments will clarify the issue. The present results also add to the mounting evidence that TSPO/MBR ligands could be used to treat fear disorders, including panic disorder, in human populations. The degree of conservation of *tspo1* indicates that anamniotes may be used as experimental models to study potential anti-panic drugs to contribute to develop of pharmacological therapeutic strategies to ameliorate panic disorders symptoms in the human.

## Acknowledgments

This manuscript first appeared as a preprint on bioRxiv (https://doi.org/10.1101/056507).

